# Identification and quantification of alternative polyadenylation sites in single cell RNA-seq data using scPAISO

**DOI:** 10.1101/2025.08.20.669565

**Authors:** Yongjie Liu, Peiwen Xiong, Songyang Li, Xinjia Liu, Tao Liu, Qinglan Yang, Shuting Wu, Hongyan Peng, Yana Li, Lingling Zhang, Yafei Deng, Yong Zhu, Junping Wang, Youcai Deng

## Abstract

Alternative polyadenylation (APA) is a critical posttranscriptional mechanism that generates transcriptomic diversity through the production of mRNA isoforms with distinct 3’ UTRs or coding sequences. Current APA analysis based on single-cell RNA sequencing (scRNA-seq) for establishing cell type-specific APA landscapes primarily rely on Read2 data, which lacks precise cleavage site (CS) information. This limitation restricts their ability to achieve precise *de novo* mapping of polyadenylation sites (PASs). Here, we present **s**ingle-**c**ell **P**oly**A**denylation **ISO**form quantification (scPAISO), a computational pipeline designed for *de novo* identification of PAS and quantification of PAS isoforms in scRNA-seq data, by leveraging the often-discarded Read1 from 3’ tag-based scRNA-seq protocols. Unlike existing tools, scPAISO directly captures mRNA 3’ end cleavage sites, enabling superior performance in motif enrichment (stronger AAUAAA signal) and peak precision (sharper PAS peaks). Moreover, the smaller peak widths enhance the spatial resolution, enabling more accurate detection of closely spaced PASs in the genome. By integrating Read1 and Read2 data, scPAISO achieves isoform-level quantification with an assignment accuracy exceeding 95%. We demonstrate the robustness of scPAISO in identifying PASs and quantifying APA events across diverse biological contexts, including hematopoiesis, systemic sclerosis (SSc), and mouse tissues. We identified stage-specific 3’ UTR lengthening in hematopoietic progenitors, global 3’ UTR remodeling in SSc and tissue-specific polyadenylation (PA) preference along with RNA-binding proteins in mice. Overall, scPAISO represents a significant advancement in the analysis of APA at single-cell resolution and provides a powerful tool for exploring the regulatory landscape of APA, offering new insights into transcriptome complexity and gene regulation in both health and disease.

## Introduction

The lengths of mRNAs and their 3’ untranslated regions (UTRs) are determined by pre-mRNA cleavage and polyadenylation, processes initiated when polyadenylation site (PAS)-regulating proteins recognize the PAS signal^1^. Most mammalian genes contain multiple PASs, enabling a single gene to produce diverse mRNA isoforms through alternative polyadenylation (APA), thereby significantly contributing to transcriptomic diversity and complexity^2^. In humans, approximately half of all genes utilize different PASs—either proximal or distal PASs within the same 3’ UTR—to generate mRNA isoforms that encode the same protein^3^. Additionally, some genes employ intronic or exonic polyadenylation (iPA or ePA) signals, producing mRNA isoforms that yield distinct protein isoforms^3^. The resulting diverse 3’ ends affect various stages of the RNA life cycle. For instance, variations in 3’ UTR length can alter the availability of binding sites for microRNAs and RNA-binding proteins, thereby regulating mRNA stability, localization, and translation^4,5^. Truncated coding or noncoding transcripts may arise through APA at intronic or exonic sites^6,7^. APA is tightly regulated, with tissue-specific, cell type-specific, or state-specific patterns^8–11^. APA occurs within diverse biological contexts, including cell differentiation^12,13^, cell proliferation^14^, embryonic development^8^, and immune responses^15,16^, and is dysregulated in diseases^11,17^.

Existing techniques for direct APA measurement, such as 3’-seq^18^, 3P-seq^19^, PAS-seq^20^, and poly(A)-seq^21^, rely on bulk RNA isolation, yielding only averaged PAS profiles across heterogeneous cell populations. In contrast, single-cell RNA sequencing (scRNA-seq) protocols, such as the 10X Chromium system^22^, capture mRNA 3’ ends via poly(A) priming, thus enabling APA analysis at single-cell resolution. Existing methods (e.g., scAPA^23^, scAPAtrap^24^, Sierra^25^, scDaPars^26^, movAPA^27^, and SCAPTURE^28^) analyze only Read2, which rarely spans mRNA 3’ end cleavage sites (CSs). Therefore, these methods serve as predictive tools rather than enabling precise PAS mapping. Although Read1 contains CS information, it is typically discarded after barcode/UMI extraction.

Here we present **s**ingle **c**ell **P**oly**A**denylation **ISO**form quantification (**scPAISO**), a pipeline that leverages both Read1 and Read2 from 3’ tag-based scRNA-seq for *de novo* mapping and quantification of PASs. scPAISO precisely identifies mRNA 3’ ends from Read1, enabling both known and novel PAS detection (half of detected PASs span <20 bp; 95% <70 bp). By anchoring Read2 to PAS peaks, scPAISO achieves isoform-level quantification, marking a major advancement in single-cell APA analysis.

## Results

### Overview of scPAISO

In the 10X Chromium 3’ single-cell RNA-seq libraries, the fragment sizes ranged from 300–600 bp. Under PE150 sequencing, Read2 usually contains a 150 bp cDNA sequence, whereas Read1 comprises a 16 bp barcode, 10/12 bp UMI, 30 bp polyT capturer sequence and ∼90 bp cDNA^22^ (**Fig. 1a, top**). Typically, only Read2 is subjected to transcriptome mapping for gene-level quantification, whereas Read1—despite containing CS information—is discarded after barcode/UMI extraction. To assess the utility of Read1 for PAS detection, we analyzed the GSE196676 dataset^29^, and mapped both Read1 and Read2 to the genome. For example, in the genomic region chr1:2,401,846-2,405,839, *RER1* is located on the sense strand and encompasses four PASs (red), whereas *PEX10* is located on the antisense strand and has two PASs (blue) located in the 3’ UTR, and all of those PASs are supported by the polyA_DB database^30^ (**Fig. 1a, bottom**). Additionally, the 3’ end of Read2 is broadly distributed, whereas the 5’ end of Read1 (CS position) presents sharper peaks (**Fig. 1a, bottom**). Moreover, two closely spaced proximal PASs of *RER1* are clearly distinguished by the Read1 signal, whereas only a single peak is observed in the Read2 signal (**Fig. 1a, bottom**). Compared with the Read2 3’ end, the transcriptome-wide Read1 5’ end is markedly enriched at transcription end sites (TESs), confirming Read1’s superiority for *de novo* PAS mapping (**Fig. 1b**).

**Fig. 1:**
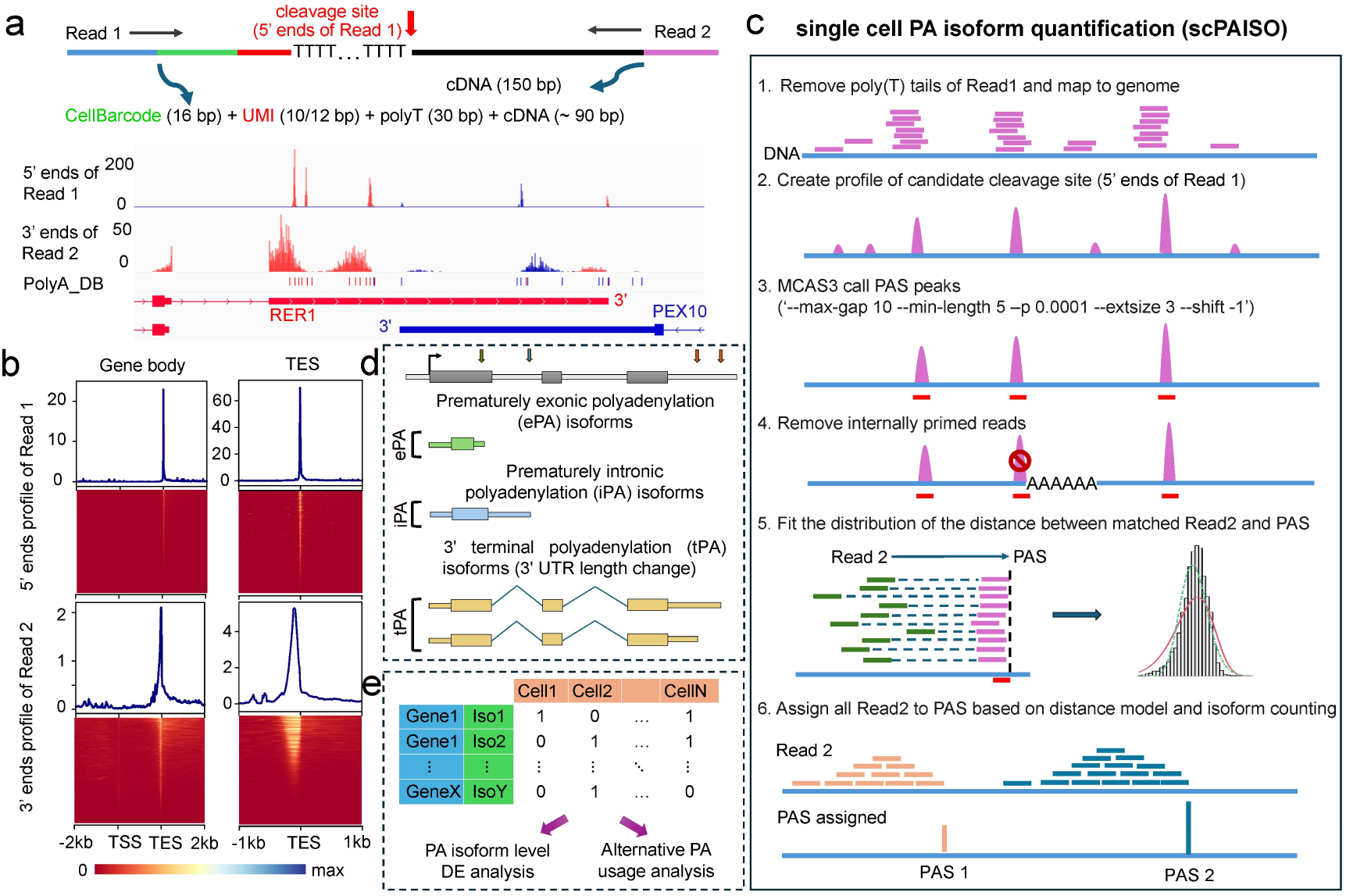
Overview of the single cell polyadenylation isoform quantification (scPAISO) pipeline for *de novo* identification and quantification of polyadenylation sites (PASs). (**a**) Schematic representation of the 10X Chromium 3’ single-cell RNA-seq library structure. Read1 contains a barcode, a UMI, a polyT capture sequence, and cDNA, and Read2 consists of a 150 bp cDNA sequence. The sharp peak at the end of Read1 indicates the precise cleavage site (CS) of the *RER1* and *PEX10* mRNA 3’ ends, enabling accurate PAS identification. Genes on the sense strand and their reads are marked in red, while those on the antisense strand are marked in blue. (**b**) Comparison of Read1 and Read2 enrichment at gene bodies and transcription end sites (TESs). (**c**) Workflow of the scPAISO pipeline, including Read1 alignment, PAS identification, model fitting, and PA isoform quantification. (**d**) Classification of polyadenylation (PA) isoforms into three main categories: exonic polyadenylation (ePA), intronic polyadenylation (iPA), and terminal polyadenylation (tPA). (**e**) Application of scPAISO for differential expression analysis and relative APA usage at single-cell resolution.

The scPAISO pipeline comprises the following six steps to systematically identify PASs and quantify PAS usage: (1) STAR-based genome alignment of Read1; (2) extraction of the Read1 5’ end position to generate cleavage site profiles; (3) candidate PAS peak calling using MACS3 based on the above profile^31^; (4) filtering of internal priming artifacts (6-mers ‘AAAAAA’ within-5/+20 bp of PAS) ^8^; (5) extraction of mapped Read1–Read2 pairs to construct a positive control dataset and fit the distribution of the distance between matched Read2 and the corresponding PAS; and (6) assign all Read2 to PASs based on the distance model and PA isoform counting (**Fig. 1c** and **Methods**).

Based on the position of PASs in transcripts, PA isoforms are divided into three categories: prematurely exonic polyadenylation (ePA) isoforms, prematurely intronic polyadenylation (iPA) isoforms and 3’ terminal polyadenylation (tPA) isoforms **(Fig. 1d**). The resulting single-cell PA isoform quantitative matrix was used for differential expression analysis of PA isoforms and APA analysis (**Fig. 1e**). A single gene may generate multiple PA isoforms, and while differential expression may not be apparent at the gene level, PA isoform-level analyses may reveal significant differences. Consequently, single-cell PA isoform-level quantification provides a more detailed perspective for analyzing differences in cellular expression profiles.

### scPAISO enables *de novo* identification of polyadenylation sites with high resolution in hematopoiesis

To validate scPAISO, we analyzed 10X Chromium scRNA-seq data from 48,500 bone marrow cells (GSE196676), including approximately 27,000 cells from healthy donors and ∼21,500 cells from patients with immune thrombocytopenia (ITP)^29^. Via graph-based Leiden clustering, and CellTypist annotation^32^, 22 cell populations, such as hematopoietic stem cells (HSC), multipotent progenitors (MPPs), dendritic cells (DCs), and six types of B cell populations, were identified (**Fig. 2a** and **Supplementary Fig. 1a**). scPAISO identified 30,626 sharp PASs (95% < 69 bp), of which 64.6% overlapped with entries in the public polyA_DB database^30^, and 68.5% were located within 500 bp of annotated PASs. For example, we identified three distinct tPAs within the 1,047 bp 3ʹ UTR of *TCF19*, a pair of antisense overlapping tPAs in the extended 3’ UTR of *TESK1* and the 3’ UTR of *CD72*, and two distinct tPAs within the 278 bp 3ʹ UTR of *SNRPG*. Despite the short 3’ UTR of *SNRPG*, Read1 signals allowed the identification of two closely spaced yet clearly distinguishable PASs, whereas Read2 captured only a single peak (**Fig. 2b**). The average 5’ end signal profiles of Read1 and 3’ end signal profiles of Read2 at the PASs further demonstrated that Read1, but not Read2, can be used for accurate detection of PASs (**Fig. 2c**). Among all the genes with predicted PASs, 56% contained alternative PASs (**Fig. 2d**). The majority of PASs were distributed in the 3’ UTR or last exon, with 374 (1.22%) PAS located in extended 3’ UTRs. Additionally, approximately 25.05% of the PASs were in the introns (**Fig. 2e**). Further enrichment analysis revealed a corresponding AAUAAA signal 50 bp upstream of the PAS^27^ (**Fig. 2f-g and Supplementary Fig. 1b**). Most annotated PASa contain the AATAAA motif or its single-nucleotide variants, except for PASs located in other exons (**Supplementary Fig. 1b**).

**Fig. 2:**
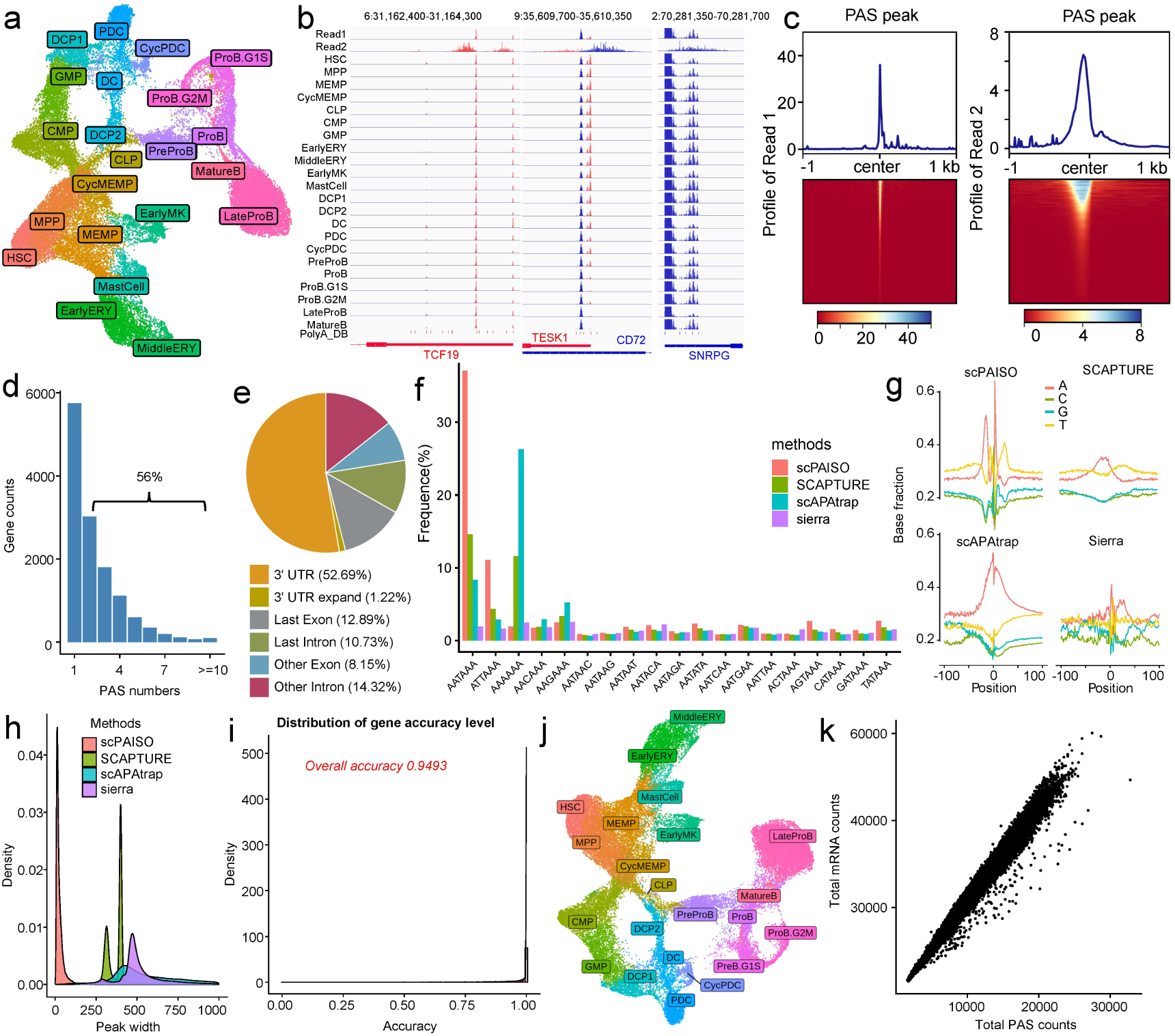
scPAISO identifies and quantifies PASs in bone marrow hematopoiesis. (**a**) UMAP visualization of scRNA-seq data from bone marrow cells. (**b**) Examples of identified PASs in the 3’ UTR of *TCF19*, antisense overlapping PASs in *TESK1* and *CD72*, and closely spaced PASs in the 3’ UTR of *SNRPG*. The IGV track was set to ‘autoscale’ for the *TCF19* and *TESK1* loci, which autoscaled selected tracks to their groupwise maximum. The IGV track maximum was set to 10 for the *SNRPG* locus. (**c**) Average Read1 and Read2 signal profiles at identified PAS. (**d**) Number of genes with alternative PAS usage. (**e**) Genomic distribution of identified PASs. (**f**) Proportional distribution of PASs containing the AATAAA motif or its 1-nt variants identified by different methods. (**g**) Nucleotide composition surrounding PASs is identified by different methods. (**h**) Peak width distribution of PASs identified by different methods. (**i**) Accuracy of PAS prediction using the *gamma distribution* model. (**j**) UMAP visualization of cells based on single-cell PA isoform expression, showing high consistency with gene-level clustering. (**k**) Correlations between gene expression matrices from Cell Ranger and scPAISO. Abbreviations: HSC, hematopoietic stem cell; MPP, multipotent progenitor; MEMP, myeloid-erythroid progenitor cell; CycMEMP, cycling MEMP; CMP, common myeloid progenitor; GMP, granulocyte-monocyte progenitor; CLP, lymphoid progenitor cell; CDP, DC progenitor; DC, dendritic cells; PDC, plasmacytoid dendritic cells; ERY, erythroid cell, PreProB, preprogenitor B cell, ProB, progenitor B cell, LateProB, late progenitor B cell.

We next compared the reliability of PAS identification between scPAISO and three Read2-based methods (SCAPTURE^28^, scAPAtrap^24^, and Sierra^25^). PASs identified by scPAISO (30,626 PASs) represented the greatest proportion of the AATAAA motif, followed by those identified by SCAPTURE (36,205 PASs) (**Fig. 2f**). In contrast, methods such as Sierra (127,842 PASs) and scAPAtrap (193,028 PASs) detected many more PASs, including a higher proportion of noncanonical variants, suggesting a trade-off between sensitivity and motif precision. The nucleotide composition of sequences surrounding scPAISO-identified PASs was also more consistent with that reported in previous studies^33^ (**Fig. 2g**). To assess the resolution of PAS identification, we compared the distribution of peak widths across different methods. Compared with the other methods, scPAISO yielded the smallest peak widths, indicating higher precision in terms of PAS detection (**Fig. 2h)**. Narrower peaks facilitate the detection of closely spaced PASs, such as the PASs of *RER1* (**Fig. 1a**) and *SNRPG* (**Fig. 2b**). We further compared the distance distributions between the PASs and their adjacent counterparts across different methods and reported that scPAISO could more effectively distinguish adjacent PASs (**Supplementary Fig. 1c**).

We then fitted three different distributions to model the distances from the 3’ ends of Read2 to the PASs in the positive control dataset and used these fitting models to predict the most likely PASs for all Read2 sequences. Among these distributions, the *gamma distribution* had the best fitting effect, with an overall accuracy of 94.9% (**Fig. 2i** and **Supplementary Fig. 1d-e**). Via the use of this model, we predicted PA isoform expression in each single cell. In some previous studies, Read2 sequences were simply assigned to its closest PAS^8,34^, whereas in our positive control dataset, approximately 11% of Read2 sequences contained a PAS that was not the closest (**Supplementary Fig. 1f**). Overall, PA isoforms located in the 3’ UTR tended to have higher expression levels, whereas those located in introns typically presented lower expression levels (**Supplementary Fig. 1g**). The proportion of PA isoforms located in the 3’ UTR and last exon was greater than those of the other regions (**Supplementary Fig. 1h**).

Dimensionality reduction and clustering analysis based on single-cell PA isoform expression levels revealed high consistency between gene and PA isoform-level UMAP visualization structures (**Fig. 2j**). Further analysis revealed a strong correlation between Cell Ranger’s gene expression matrix and the merged gene expression matrix of the PA isoforms, indicating that the results of the two methods were consistent (**Fig. 2k**). These results demonstrate that scPAISO not only accurately identifies PASs but also quantitatively measures PA isoform levels at the single-cell level.

### scPAISO enables the analysis of single-cell PA isoform dynamics during hematopoiesis and ITP-associated APA reprogramming

Compared with conventional mRNA quantification, single-cell PA isoform profiling offers a higher-resolution lens for dissecting cellular subpopulations. We used the FindMarkers function from the Seurat package^35^ to identify differentially expressed genes or PA isoforms (adjusted p-value < 0.01 and |avg_log2FC| > 0.3) during hematopoiesis. For some genes, although no significant changes were observed at the gene level, PA isoform alterations were detected during hematopoiesis (**Fig. 3a**). During the differentiation of HSCs into MPPs (HSC→MPP), 354 PA isoforms underwent specific changes, most of which were upregulated. For these ISO-only genes, a consistent trend (upregulated or downregulated) was observed between gene-level changes and PA isoform-level changes; however, the magnitude of changes at the gene level was relatively small (**Fig. 3b**). During the differentiation of progenitor B cells (ProB) into late progenitor B cells (LateProB) (ProB→LateProB), 2,145 PA isoforms significantly changed, most of which were downregulated. GO enrichment analysis revealed that ISO-only PA isoforms involved in the HSC→MPP transition were associated with histone modification, stem cell population maintenance, positive regulation of myeloid cell differentiation, and other processes (**Fig. 3c left**). ISO-only PA isoforms in ProB.G1→LateProB transitions were associated with DNA-templated transcription initiation, RNA methylation, TOR signaling, telomere maintenance, the metaphase/anaphase transition of mitotic cell cycle, and B-cell activation involved in immune responses (**Fig. 3c right**). For example, the expression of histone acetyltransferase 1 (*HAT1*), a gene that enhances proliferation by coordinating histone production, acetylation, and glucose metabolism^36^, did not significantly differ between HSCs and MPPs. However, one of its tPA isoforms was significantly upregulated in MPPs (**Fig. 3d top**). T-cell acute lymphocytic leukemia 1 (*TAL1*), which regulates immature human hematopoietic cell self-renewal^37^, also showed no gene-level differential expression between HSCs and MPPs, but one of its iPAs was significantly downregulated in MPPs (**Supplementary Fig. 2a**). *TEN1*, a gene essential for telomeric C-strand synthesis^38^ exhibited no differential gene expression, but its two nearby tPAs (∼120 bp apart) were highly expressed in proB cells, suggesting a potential role in telomere maintenance (**Fig. 3d bottom**). *WDR33*, which encodes a key component of the mRNA polyadenylation machinery^39^, also increased the expression of two tPAs in proB cells, suggesting its involvement in 3’ UTR length regulation during B cell development (**Supplementary Fig. 2b**).

**Fig. 3:**
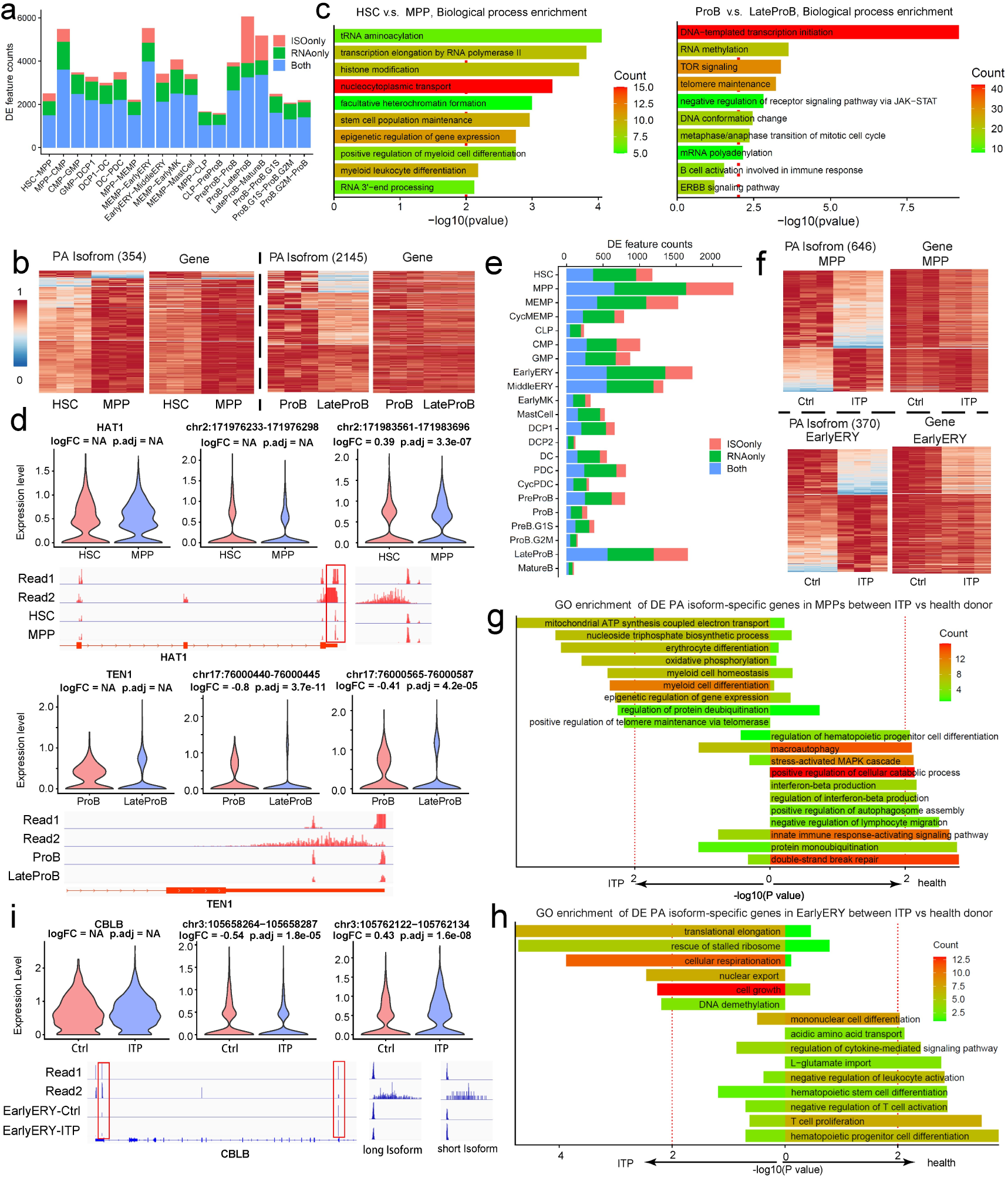
PA isoform-specific differential expression. (**a**) Number of genes or PA isoforms that are differentially expressed during hematopoiesis. (**b**) Heatmap showing specific changes in PA isoform expression during HSC-to-MPP differentiation and ProB-to-LateProB transition. *ISOonly*: differential expressions were detected only at the PA isoform level. *RNAonly*: differential expressions were detected only at the gene level. *Both*: differential expressions were detected at both the gene and PA isoform levels. (**c**) Gene Ontology (GO) enrichment analysis of specific differentially expressed PA isoforms during HSC differentiation (left) and B cell development (right). (**d**) *HAT1* (above) and *TEN1* (bottom) showing PA isoform-specific changes without corresponding gene-level expression differences. Read1 track: 5’ ends profile of Read1 from all cells. Read2 track: 3’ ends profile of Read2 from all cells. Cell type track: a predicted 3’ ends profile of Read1 from given cells. (**e**) Number of differentially expressed genes or PA isoforms in ITP. (**f**) Heatmap showing specific changes in PA isoform expression in MPPs (top) and EarlyERYs (bottom). (**g-h**) GO enrichment of upregulated or downregulated PA isoforms in MPPs (**g**) and EarlyERYs (**h**) between normal and ITP cells. (**i**) The expression of CBLB did not significantly differ at the total RNA level in EarlyERYs, but its short isoform was highly expressed in ITP, while the long isoform was expressed at reduced levels.

Even broader PA isoform-specific expression changes were observed in each subset between healthy donor and ITP samples (**Fig. 3e**). There were 646 differentially expressed PA isoforms in MPPs and 370 differentially expressed PA isoforms in early erythrocytes (EarlyERYs) (**Fig. 3f**). Upregulated PA isoforms in MPPs from patients with ITP were associated with processes such as mitochondrial ATP synthesis coupled electron transport, erythrocyte differentiation, myeloid cell differentiation, and oxidative phosphorylation, whereas downregulated PA isoforms in MPPs from patients with ITP were associated with processes such as innate immune response-regulating signaling pathway, interferon-beta production and regulation of hematopoietic progenitor cell differentiation (**Fig. 3g**). For example, BRD1 is required for the transcriptional activation of erythroid developmental regulator genes^40^, whose iPA is upregulated in MPPs from patients with ITP (**Supplementary Fig. 2c**). In addition, upregulated PA isoforms in EarlyERYs from patients with ITP were linked to translational elongation and rescue of stalled ribosome, which are closely linked to the hallmark biological processes of early erythropoiesis—ribosome biogenesis and hemoglobin accumulation^41^; whereas downregulated PA isoforms in EarlyERYs from patients with ITP were associated with hematopoietic progenitor cell differentiation, T cell proliferation, negative regulation of T cell activation, and regulation of cytokine-mediated signaling pathways (**Fig. 3h**). Cbl proto-oncogene B (CBLB) is associated with the biological process of T cell proliferation and it acts as a critical negative regulator of JAK2 signaling in erythroid cells by cooperating with LNK to promote JAK2 degradation and restrain cytokine-induced proliferation^42^; however, its role in the marked increase of EarlyERYs observed in patients with ITP remains unclear, and no significant difference in *CBLB* expression was detected at the total RNA level in EarlyERYs. Through differential expression analysis at PA isoform level, we determined that the short isoform of *CBLB* was upregulated in ITP, whereas the long isoform downregulated in ITP (**Fig. 3i**). This suggests that the switch in PA isoforms may increase the proliferation of EarlyERY by reducing the levels of full-length CBLB protein. Eukaryotic translation initiation factor 4E type 2 (EIF4E2) is involved in the regulation of translation^43^, and the ePA of *EIF4E2* is upregulated in EarlyERYs in ITP (**Supplementary Fig. 2d**). SOS1 is a guanine nucleotide exchange factor for Ras proteins and plays a crucial role in the immune response^44^, and its tPA level was significantly was downregulated in EarlyERYs from patients with ITP (**Supplementary Fig. 2e**).

In summary, our analysis demonstrated that scPAISO achieves highly accurate quantification at the PA isoform level. Compared with conventional gene-level quantification, this method provides superior resolution in expression profiling, enabling the detection of PA isoform-specific differential expression patterns. This enhanced granularity will significantly facilitate more precise functional genomic studies.

### Nucleotide-resolution tracking of dynamic PAS selection during hematopoietic differentiation

To characterize global patterns of 3’ UTR dynamics during hematopoietic differentiation, we analyzed the above-mentioned bone marrow scRNA-seq data (GSE196676) using PASTA, a computational framework for single-cell PAS heterogeneity analysis^34^. First, we processed the PAS count matrices using Dirichlet multinomial regression to quantify the residual variability of the PA isoforms, which characterized the heterogeneity in the relative use of PASs at single-cell resolution.

Dimension reduction was performed directly on the polyA residuals, identifying seven distinct polyA clusters (resolution=0.4). We observed less cell type-specific specificity in polyA clusters (**Fig. 4a-b**).

**Fig. 4:**
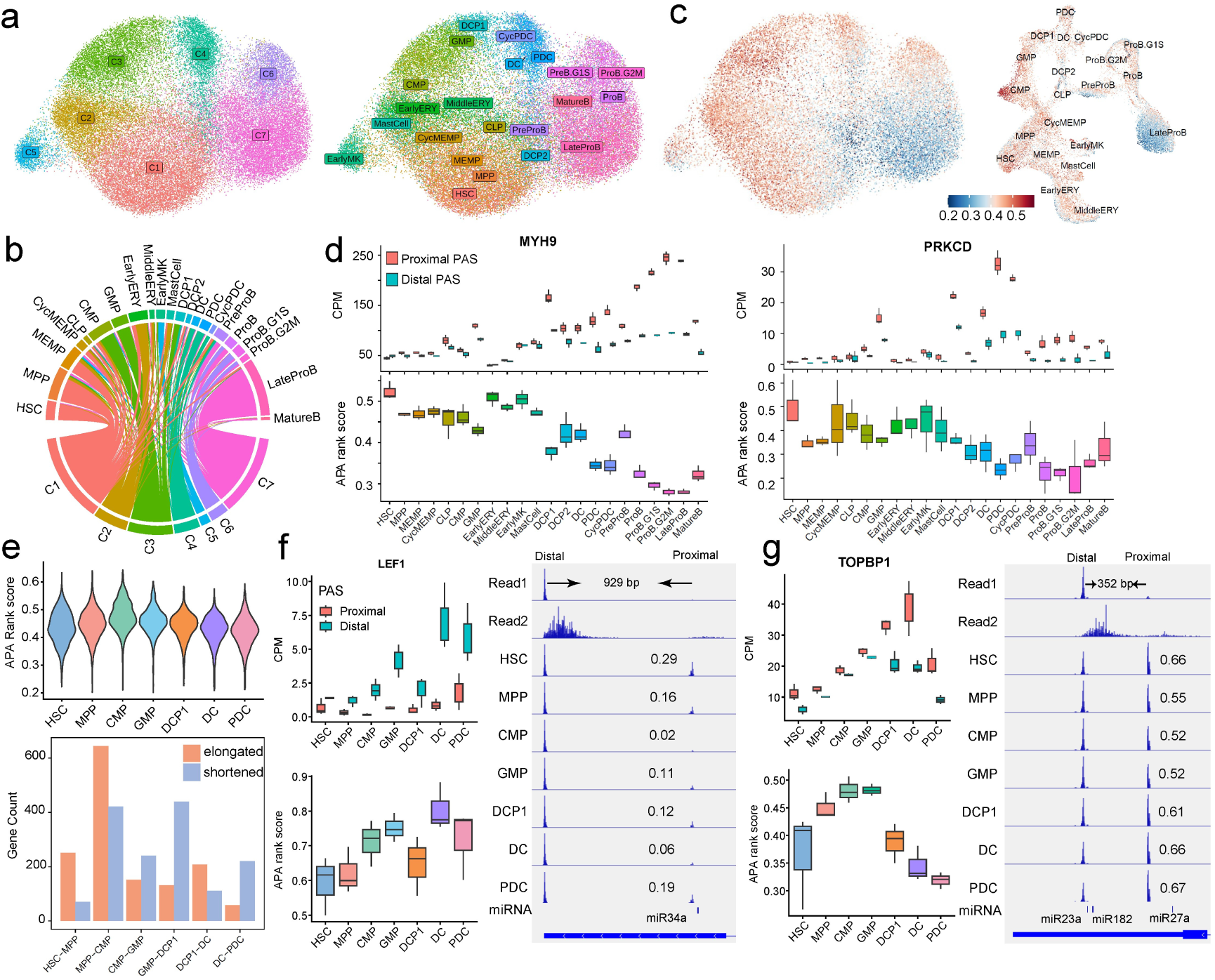
Dynamic changes in 3’ UTR length during hematopoiesis. (**a**) UMAP visualization of PA clusters (left) and cell annotation (right) based on PAS usage residuals. (**b**) Matching relationships of cell types between PA usage residual-based clusters and gene-based cell annotation. (**c**) APA rank scores for relative PAS usage across different cell types. Viewed by PAS usage based UMAP (left) and gene expression based UMAP (right). (**d**) CPM (counts per million)-normalized PA isoform expression and APA rank scores of *MYH9* (left) and *PRKCD* (right). (**e**) APA rank score trends during DC differentiation (top). Gene numbers with elongated or shortened 3’ UTR lengths (bottom). (**f-g**) CPM normalized PA isoform expression and PAS preference of *LEF1* (**f**) and *TOPBP1* (**g**) during DC differentiation. To illustrate changes in the PAS usage ratios, the track in the IGV plot is set to “autoscale”. The utilization ratio of the proximal PAS was explicitly labeled in the diagram. The distance between two adjacent PASs is labeled.

We next developed a rank score (ranging from 0–1) to quantify preferential PAS selection for each gene and evaluate the PAS usage bias across hematopoietic lineages. A rank score of 0 indicates exclusive proximal PAS, while a score of 1 indicates exclusive distal PAS. C2 and C3 clusters which had the highest rank scores, included common myeloid progenitors (CMPs), granulocyte–monocyte progenitors (GMPs), DC progenitor 1 (DCP1), EarlyERYs and middle erythroid cells (MiddleERYs), suggesting preferential distal PAS usage (**Fig. 4b-c**). The C7 cluster (lowest rank score) included ProB and LateProB cells, reflecting proximal PAS bias (**Fig. 4b-c**).

Next, we systematically identified genes with PAS relative usage dynamics during hematopoiesis via the PASTA package’s FindDifferentialPolyA function (Bonferroni-corrected p< 0.05, Δusage> 0.05). A total of 6,121 PA isoforms corresponding to 2,001 genes with stage-specific PAS switching were identified. To investigate 3’ UTR length dynamics during hematopoiesis, we aggregated single-cell data by cell type to generate a pseudobulk polyadenylation (PA) quantification matrix. Using this matrix, we computed gene-specific rank scores reflecting PAS usage preference across cell types and performed zscore normalization to standardize the data. We subsequently applied k-means clustering (k=10) to identify distinct patterns of the 3’ UTR length, and we revealed dynamic 3’ UTR length changes throughout hematopoietic differentiation, with stage-specific selection of PASs (**Supplementary Fig. 3a**). Several key regulatory genes exhibited dynamic 3’ UTR lengthening/shortening patterns during hematopoietic differentiation. For example: Myosin heavy chain 9 (*MYH9*), which critically regulates HSPC survival^45^, and B cell activation and proliferation^46^, exhibited longer 3’ UTRs (preferring the distal PAS) in primitive progenitors (HSCs/MPPs) but shorter 3’ UTRs (preferring the proximal PAS) in committed lineages (e.g., plasmacytoid dendritic cells (PDCs)/B cells) (**Fig. 4d left** and **Supplementary Fig. 3b**). Notably, the distal PAS was located only 256 bp downstream of the proximal PAS, and this short region contains multiple miRNA binding sites^47^ (**Supplementary Fig. 3b**). This not only demonstrates the high resolution of scPAISO in detecting closely spaced PASs but also highlights its ability to identify potential miRNA target^48^ and post-transcriptional regulatory mechanisms mediated by APA. Protein kinase C delta (*PRKCD*), a key modulator of HSPC proliferation and metabolism^49^, similarly showed a distal PAS (only 157 bp away from the proximal PAS) preference in undifferentiated HSPCs (**Fig. 4d right** and **Supplementary Fig. 3c**).

PolyA residual based clustering revealed a continuous APA usage change from HSCs (C1), MPPs (C1), CMPs (C3), GMPs (C3), DCP1 (C3), and DCs (C4) to PDCs (C4).

To investigate PAS usage dynamics during HSC differentiation into PDCs, we quantified rank scores across developmental stages. The rank scores gradually increased from HSC stage and peaked at CMP stage but then decreased later in differentiation (**Fig. 4e, top**). We then clustered genes showing significant PAS usage changes (Bonferroni-corrected p< 0.05, Δusage> 0.05) and identified the most significant changes in PAS usage occurring during the transition from MPP to CMP (**Fig. 4e, bottom**). For example, LEF1, a transcription factor for myeloid progenitor proliferation and differentiation^50^, showed greater distal PAS preference (high rank score) in CMPs/GMPs (**Fig. 4f**). To illustrate the changes in PAS usage ratios, the track scale in the IGV was set to ‘autoscale’. The utilization ratio of the proximal PAS was explicitly labeled. The proximal PAS of *TEL1* accounted for 29% and 16% in HSCs and MPPs, respectively, but accounted for only 2% and 11% in CMPs and GMPs. Similarly, TOPBP1, which encodes a key DNA topoisomerase for DNA replication/repair processes^51^, also exhibited a preference for a distal PAS (only 352 bp away from the proximal PAS) in CMPs/GMPs (**Fig. 4g**).

### PAS usage preferences in SSc

SSc is a progressive autoimmune disorder characterized by pathological collagen deposition in cutaneous and visceral tissues. Using scPAISO, we analyzed scRNA-seq data from the skin tissue of 22 patients with SSc and 18 healthy donors^52^. After stringent quality filtering (retaining 95,954 high-quality cells), we categorized the transcriptome into 12 major cell types (**Fig. 5a** and **Supplementary Fig. 4a**). scPAISO identified 33,683 sharply PAS peaks, with 95% localized within 72-bp regions. For example, *RCAN3* exhibited three specific tPAs in its 3’ UTR (**Supplementary Fig. 4b**). Nucleotide composition near the PAS aligned with the canonical motif (**Supplementary Fig. 4c**). When assigning Read2 to PAS, a Read2 assignment accuracy of 96.78% was achieved using *Weibull distribution* modeling (**Supplementary Fig. 4d**). UMAP visualization of cell-type signatures confirmed strong concordance between gene and PA-isoform-level clustering (**Fig. 5a**).

**Fig. 5:**
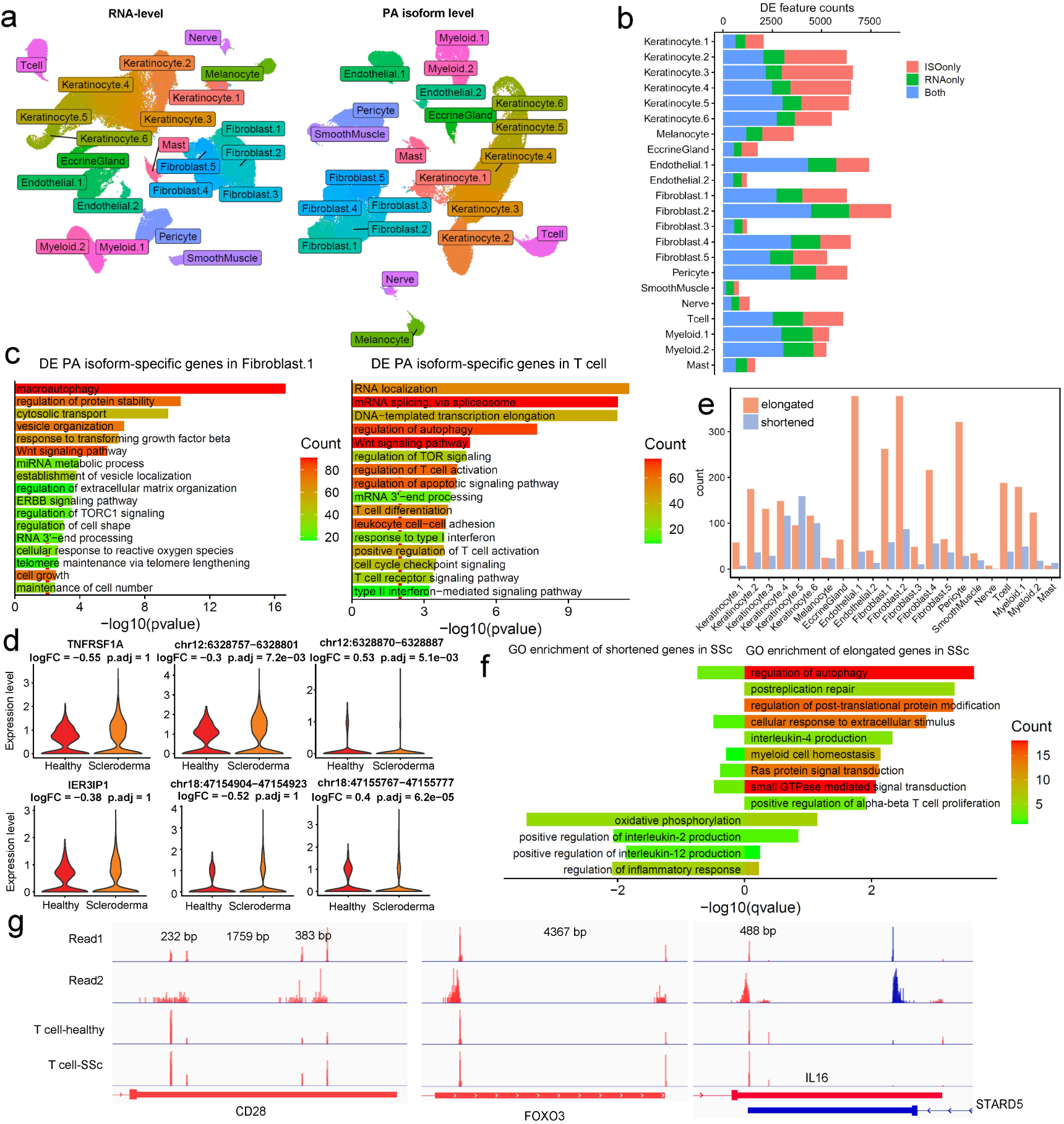
PA isoform preference in systemic sclerosis (SSc). (**a**) UMAP visualization of gene expression (left) and PA isoform (right) data from patients with SSc and healthy controls. (**b**) Number of differentially expressed genes or PA isoforms in SSc. (**c**) GO enrichment analysis of specific differentially expressed PA isoforms in Fibroblast.1 (left) and T cells (right). (**d**) *TNFRSF1A* (top) and *IER3IP1* (bottom) showing PA isoform-specific changes without corresponding gene-level expression differences. (**e**) Gene numbers with dynamic changes in 3’ UTR length between SSc and controls. (**f**) GO enrichment analysis of genes with elongated or shortened 3’ UTRs in T cells. (**g**) PAS preference changes in *CD28, FOXO3* and *IL16*. The distance between two adjacent PASs is labeled.

Disease-associated PAS usage analysis revealed widespread differential expression of PA isoforms without corresponding changes in total gene expression across most cell populations (**Fig. 5b**), highlighting posttranscriptional dysregulation. In Fibroblast.1, differentially expressed PA isoform-specific genes were implicated in macroautophagy, the ERBB signaling pathway, and regulation of TORC1 signaling (**Fig. 5c**). Notably, both IER3 interacting protein 1 (*IER3IP1*) and tumor necrosis factor receptor superfamily member 1A (*TNFRSF1A*), which are involved in the regulation of fibroblast apoptosis^53,54^, were differentially expressed PA isoform-specific genes (**Fig. 5d**). In T cells, differentially expressed PA isoform-specific genes were linked to regulation of T cell activation and differentiation, leukocyte cell–cell adhesion, and response to type I interferon (**Fig. 5c**).

To further investigate 3’ UTR length remodeling in SSc, we quantified rank scores and compared healthy donors and patients with SSc across cell populations. Strikingly, global 3’ UTR lengthening was observed in disease-associated cell types, including Fibroblast.1, T cell, and myeloid lineages (**Fig. 5e**). Functional implication analysis of 3’ UTR remodeling in T cells revealed that genes with elongated 3’ UTRs were enriched for interleukin-4 production and positive regulation of alpha-beta T cell proliferation, while genes with shortened 3’ UTRs were strongly associated with positive regulation of interleukin-2 and interleukin-12 production, and regulation of inflammatory responses (**Fig. 5f**). Key regulatory genes in T cells exhibited differential PAS usage. *CD28* (T cell costimulation and activation) and *FOXO3* (critical for T cell homeostasis and survival)^55,56^ were associated with a distal PAS preference in SSc. In contrast, *IL16*, which is involved in immune response, preferentially used the proximal PAS in SSc^57^ (**Fig. 5g**).

### PAS usage landscape across mouse tissues

Although PA dynamics during mouse embryogenesis have been explored^8^, a systematic characterization of PAS usage—particularly at single-cell resolution—remains incomplete for both the embryonic and adult stages, owing to limitations in precise PAS mapping. To address this gap, we implemented scPAISO on a comprehensive scRNA-seq dataset containing cells from brain, liver, muscle, and skin tissues and hematopoietic lineages (isolated from bone marrow, peripheral blood, and the spleen)^58^. After quality filtering (retaining 265,947 cells) and tissue-specific dimensionality reduction and annotation (**Supplementary Fig. 5a**), we integrated the data and reanalyzed the cells for unified visualization (**Fig. 6a**). Next, we applied scPAISO to this dataset and identified a total of 51,926 PASs. The nucleotide composition surrounding the 3’ UTR PASs was consistent with those in previous reports (**Supplementary Fig. 5b**). When Read2 data were assigned to PAS, the *Weibull distribution* yielded the best fit, achieving an overall accuracy of 95.91% (**Supplementary Fig. 5c**). Notably, we observed distinct tissue-specific PAS usage patterns: immune cells displayed systematically higher rank scores, suggesting a bias toward distal PAS selection, whereas parenchymal cells—particularly those from the liver, brain, and muscle—showed lower rank scores, indicating a preference of the proximal PAS (**Fig. 6a**).

**Fig. 6:**
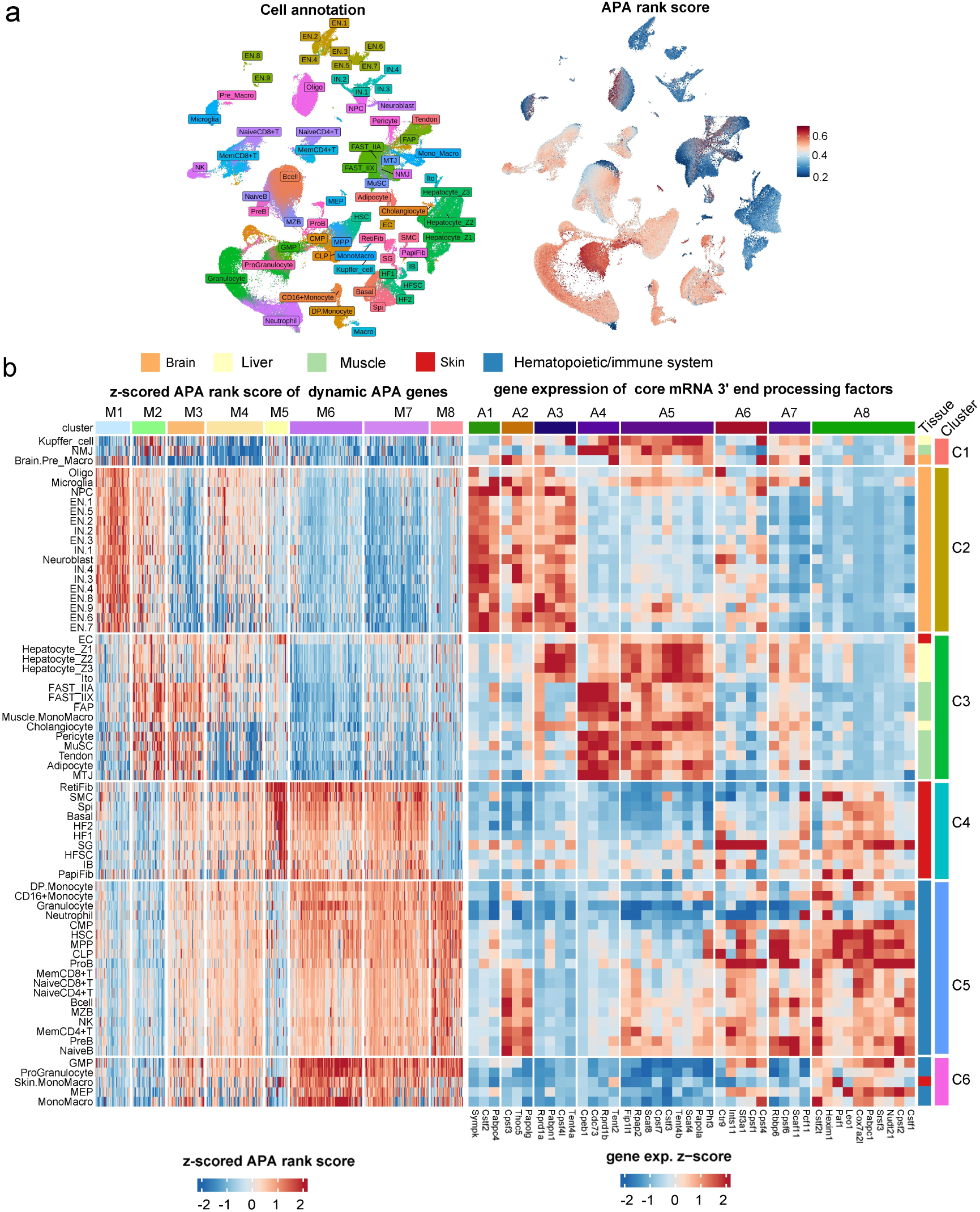
Tissue and cell-specific PAS usage preferences in mouse tissues. (**a**) UMAP visualization of cell annotation (left) and APA rank scores (right) from the scRNA-seq data of multiple mouse tissues. (**b**) Left: heatmap of APA rank score patterns of genes with significant PAS usage changes. Right: heatmap of gene expression patterns of mRNA 3’ end processing factors. Abbreviations: Brain: EN, excitatory neurons; IN, inhibitory neurons; NPC, neural progenitor cells; Oligo, oligodendrocytes. Skeletal muscle: FAP, fibro-adipogenic progenitor cells; FAST, fast-twitch muscle fibers; MTJ, myotendinous junction; NMJ, neuromuscular junction; MuSCs, muscle stem cells. Liver: Ito, Hepatic stellate cells. Skin: Basal, stratum basale keratinocyte; EC, endothelial cell; HFSCs, hair follicle stem cells; HF, hair follicle; IB, inner bulge; PapiFib, papillary fibroblast; RetiFib, reticular fibroblast; SG, sebaceous gland cells; SMC, smooth muscle cell; Spi, stratum spinosum keratinocyte. Hematopoietic/immune system: HSC, hematopoietic stem cell; MPP, multipotent progenitors; MEP, myeloid-erythroid progenitor; CMP, common myeloid progenitor; GMP, granulocyte-monocyte progenitor; CLP, lymphoid progenitor.

To further characterize PAS usage heterogeneity across diverse cell types, we identified 5,220 genes that exhibited significant variation in 3’ UTR PAS selection. Based on the result of the rank score analysis, these genes were clustered into eight modules, and the cells were segregated into six major categories (**Fig. 6b, left**). Clustering analysis revealed that immune lineages (primarily C5 and C6) and dermal parenchymal cells (primarily C4) exhibited coordinated PAS selection profiles. Despite distinct ontogeny, parenchymal cells from striated muscle and hepatocytes (C3) were clustered together with comparable PAS rank score distributions (**Fig. 6b, left**). Striking examples of cell type-specific PAS switching included: *Foxo3*, which was highly expressed in C1 and C2 but showed preferential distal PAS usage (higher rank scores) in C4 to C6, implying uncoupled regulation of transcript levels and 3’ UTR isoform selection (**Supplementary Fig. 6a**); *Pbx2* used the distal PAS from C1 to C4 but shifted to proximal PAS in C5 and C6 (**Supplementary Fig. 6b**).

To explore the transcriptional programs governing PAS selection dynamics across cell types, we profiled the expression of core mRNA 3’ end processing factors^34^ and subjected them to unbiased clustering (**Fig. 6b, right**). This analysis resolved eight gene modules (A1–A8), each of which demonstrated pronounced cell-type specificity. For example, *Cstf2* (module A1) and *Pabpn1* (module A3) were highly expressed in neural cells, *Cpeb1* (module A4) was highly expressed in C3 cells, and included liver and muscle tissue, while cleavage and polyadenylation specificity factor 2 (*Cpsf2*) and serine/arginine-rich splicing factor 3 (*Srsf3*)^59^ (module A8) were highly expressed in immune cells (**Supplementary Fig. 6c**). These findings suggest that cell type-specific PAS preferences are likely shaped by the combined expression dynamics of these regulatory modules.

### Coupling between RNA-binding protein (RBP) expression and 3’ UTR length

The extension of 3’ UTR length may lead to an increase in RBP binding sites. Therefore, it is reasonable to hypothesize that the overall elongation of 3’ UTRs in cells might be coupled with upregulation of the expression levels of corresponding RBPs. Accordingly, we performed k-means clustering on all annotated 1,644 RBP genes^60^ (except for core mRNA 3’ end processing factors) based on their expression profiles, resulting in ten distinct modules (R1 to R10). Module R2 was highly expressed in cell clusters C2 and C3 (**Fig. 7a**), and correspondingly, genes in Cluster M2 exhibited longer 3’ UTRs in these cell clusters (**Fig. 6b, left)**. Module R3 was highly expressed in cell cluster C2 (**Fig. 7a**), and genes in Cluster M1 displayed longer 3’ UTRs specifically in C2 (**Fig. 6b, left)**. Module R10 was predominantly expressed in cell clusters C4–C6 (**Fig. 7a**), and genes in clusters M6 and M7 had longer 3’ UTRs in these cell populations (**Fig. 6b**). For example, the expression of the *Exosc* family was increased in immune cell types (C5 and C6), whereas the expression levels of the *Dnah* and *Celf* gene families were increased in brain tissue-derived cells (C2) (**Fig. 7b)**. We propose that the upregulated expression of specific RBPs may be functionally linked to the elongation of target gene 3’ UTRs, suggesting a potential posttranscriptional regulatory mechanism. RBPs exhibit specific sequence preferences in terms of RNA binding^61,62^. To further confirm the coupling between RBP expression and 3’ UTR length, we collected motif information of RBPs from starBase v2.0^47^ and analyzed the enrichment of RBP motifs in 3’ UTRs across different cell types. Specifically, we downloaded the top-ranked motifs of different RBPs from various crosslinking-immunoprecipitation (CLIP) experiments in the starBase database, which yielded in 2,179 motif entries corresponding to 319 RBP genes (**Supplementary Fig. 7a**). Furthermore, MOODS-DNA^63^ was used for genome-wide motif scanning, and strand-matched motif information within 3’ UTRs was subsequently extracted for downstream analysis. We then extracted the genomic coordinates of the 3’ UTRs of different tPA isoforms, defined them as peaks and quantified the expression levels of these tPA isoforms as peak counts. These data were subsequently converted into a ChromVAR-compatible object^64^. Using the ChromVAR package, we ultimately quantified the enrichment of RBP binding motifs across different cell types.

**Fig. 7:**
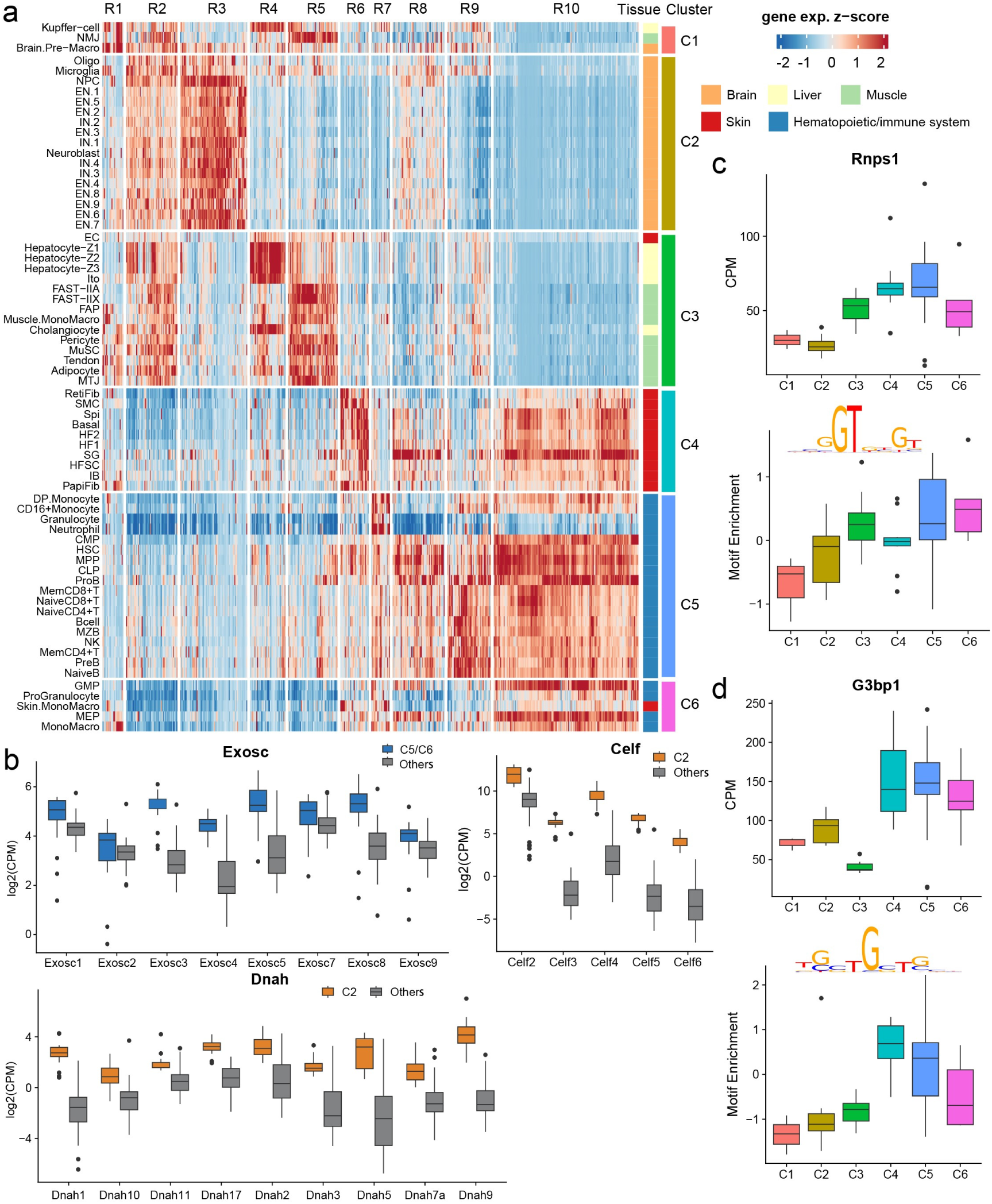
Gene expression patterns of RBP genes. (**a**) Heatmap of the gene expression patterns of RBP-related genes. (**b**) Gene expression preferences of *Exosc*-family, *Dnah*-family and *Celf*-family genes. (**c-d**) Gene expression levels and motif enrichment of *Rnps1* (**c**) and *G3bp1* (**d**) across cell clusters.

Among these RBP-motif pairs, 169 showed a positive correlation between gene expression and corresponding motif enrichment (Pearson correlation > 0.5) (**Supplementary Fig. 7b**). For example, RNA binding protein with serine repeat 1 (Rnps1), which is involved in mRNA splicing and RNA quality control, regulates hematopoietic stem cell (HSC) differentiation as well as B and T cell development and function^65^. This gene was highly expressed in cluster C5 (mainly immune cells), while its motif was also more highly enriched in the corresponding group (**Fig. 7c)**. Another example is GTPase activating protein binding protein 1 (G3bp1), which regulates mRNA stability and translation. G3bp1 plays a role in modulating inflammatory responses in macrophages and dendritic cells, and can influence antibody secretion in B cells^66,67^. It is highly expressed in clusters C4–C6, with its motif showing greater enrichment in the corresponding cell groups (**Fig. 7d**).

These results support a coordinated relationship between 3’ UTR elongation and increased expression of corresponding RBPs across cell types. The positive correlation between RBP expression and motif enrichment further suggests a potential posttranscriptional regulatory mechanism underlying 3’ UTR lengthening.

## Discussion

In this study, we present scPAISO, a novel computational pipeline designed for the *de novo* identification and quantitative analysis of PASs at single-cell resolution through 3’ tag-based scRNA-seq data. By leveraging Read1—a data source that encompasses vital 3’ cleavage site information yet is conventionally discarded in standard scRNA-seq analyses—scPAISO enables precise PAS localization and quantitation of PA isoforms within individual cells.

Previously, scRNA-seq APA pipelines inferred PAS positions from diffuse Read2 peaks, yielding ambiguous coordinates^24,25,28^. scPAISO directly utilizes Read1 to resolve PASs into sharp, discrete peaks, markedly improving positional precision, especially for intron-crossing signals and closely spaced PASs. In the *RER1* locus (**Fig. 1a**), an intron within the 3’ UTR generates a strong Read2 signal at its 5’ splice site that could be misannotated as a PAS. Read1 data confirmed that there is no internal PAS, demonstrating that the flanking reads originate from a single continuous transcript. In addition, scPAISO clearly distinguished two closely spaced proximal PASs undetectable in Read2-based analyses (**Fig. 1a**). Moreover, scPAISO PAS calls align precisely with polyA_DB^30^ annotations, underscoring the method’s fidelity and clarity (**Fig. 1a**).

Furthermore, scPAISO outperforms prior methods by identifying significantly more *de novo* PASs and enabling quantitative tracking of PA isoforms neglected by indirect inference tools. In quantifying PA isoforms, scPAISO attained >95% allocation accuracy and yielded PAS peaks that were markedly sharper (95% width <70 bp) than those produced by Read2-based tools^24,25,28^. This narrow peak profile enables reliable discrimination of adjacent PASs and resolves genes with abbreviated 3′ UTRs. For example, in *CD28* 3’ UTR (**Fig. 5i**), Read2 showed a dense, ambiguous cluster of signals, whereas Read1 resolved two discrete high-confidence PASs. A similar resolution is achieved for *TAL1*’s iPAS (**Fig. s2a**). Reexamination of these previously discarded Read1 signals markedly enriched canonical CPSF-complex binding motifs^68^: the frequencies of AATAAA and ATTAAA increased sharply around scPAISO-defined PASs compared with those around PASs obtained via other tools^24,25,28^ (**Fig. 2f**). Concomitantly, artifactual runs of adenosines, often misassigned as PASs by the impact of internal primer dimers, were significantly reduced.

Applications of scPAISO in hematopoiesis, SSc, and mouse multitissue atlases revealed unannotated PASs^30^ in genes such as *TAL1*^37^, *MYH9*^45^, *PRKCD*^49^ (hematopoiesis), and *SOS1*^44^*, CD28*^55^, and *FOXO3*^56^ (immune response and activation). scPAISO applied to multiple mouse tissues also revealed cell type-and tissue-specific preferences for PAS usage (e.g., immune cells with a distal PAS preference, parenchymal cells in brain and liver with a proximal PAS preference) (**Fig. 6b**). The expanded PASs obliges us to reassess how APA impacts the expression and activity of these pivotal genes. The selective use of these PASs can (i) reposition miRNA and RBP-binding sites or alter epigenetic marks within the affected 3′ UTR (tPA) or (ii) generate distinct protein isoforms (iPA and ePA)^3^.

Beyond the direct posttranscriptional effects of APA on gene output, specific APA signatures can serve as orthogonal biomarkers for cell-state classification during the initial clustering step of single-cell workflows. PA isoform analysis reveals expression differences that are not detectable by gene-level analysis alone. These APA-only differences recapitulate both lineage relationships and disease-associated alterations, underscoring the complementary power of single-cell APA profiling. CBLB acts as a critical negative regulator of EarlyERY proliferation^42^. Although no significant difference in the expression of *CBLB* was detected at the total RNA level in EarlyERYs, its short isoform was highly expressed in ITP, whereas the long isoform showed reduced expression in ITP, suggesting that the switch in PA isoforms may enhance the proliferation of EarlyERYs by reducing the levels of the full-length CBLB protein (**Fig. 3i**). In SSc, high-resolution PAS mapping and precise quantification by scPAISO also revealed that APA dynamics could refine disease subtyping with clinical relevance. Moreover, integrating PA-isoform dynamics into clustering algorithms thus offers an extra dimension to refine cell-type annotation and to chart subtle developmental transitions that would otherwise remain invisible.

In conclusion, in scPAISO, discarded Read1 data are repurposed to deliver single-cell PAS atlases with nucleotide resolution, uncovering APA’s roles in developmental regulation (e.g., hematopoiesis), disease mechanisms (e.g., SSc stratification), and therapeutic targeting (e.g., isoform-specific interventions). As single-cell technologies continue to evolve, tools such as scPAISO will be helpful for unraveling the complexity of the transcriptome and its regulation at the single-cell level. Moreover, current studies on 3’ UTR alternative polyadenylation quantitative trait loci (3’aQTLs) are all based on bulk RNA-seq data^69^, which inherently have significant limitations in calculating PAS usage. The development of our method could facilitate research on 3’aQTLs that is based on published scRNA-seq data^70–72^.

## Methods

### Processing of scRNA-seq data

Single-cell RNA-seq raw fastq data were obtained from publicly available datasets, including bone marrow hematopoiesis (GEO, GSE196676)^29^, SSc (GEO, GSE249279)^52^, and mouse tissues (GSA, CRA004660)^58^. The single-cell expression matrix was generated using Cell Ranger (https://github.com/10XGenomics/cellranger; GRCh38 for human, GRCm38 for mouse) and processed with the Seurat package (v4.0.1)^35^.

In hematopoiesis, low-quality cells with fewer than 2000 genes detected or more than 15% mitochondrial reads were excluded. Genes expressed in fewer than 10 cells were also removed. The remaining cells were normalized using the LogNormalize method with a scale factor of 10,000. The top 3,000 highly variable genes were selected for downstream analysis. Principal component analysis (PCA) was performed on the scaled data, and the top 30 principal components were used as inputs for Harmony (v0.1.0) integration^73^. The top 20 integrated embeddings from Harmony were then used for subsequent clustering and visualization. Cell clusters were identified at resolution = 0.8. Clusters with < 500 cells were removed. CellTypist (v1.6.3)^32^ was used for cell annotation with default parameters and was validated manually against known marker genes.

In SSc, low-quality cells with < 500 UMIs, > 50000 UMIs, > 40% ribosome reads, or > 15% mitochondrial reads were excluded. Genes expressed in < 10 cells were also removed. The top 2,000 highly variable genes were selected for downstream analysis. PCA was performed on the scaled data, and the top 50 principal components were used as inputs for Harmony (v0.1.0) integration. The top 20 integrated embeddings were employed for subsequent clustering and visualization. Cell clusters were identified at resolution = 0.6; clusters < 500 cells were removed for further analysis. Cell annotation was performed based on marker genes from prior studies.

In mouse tissues, low-quality cells with < 500 UMIs detected, > 40000 UMIs, < 200 genes, > 6000 genes, > 40% ribosome reads, or > 15% mitochondrial reads were excluded. Genes expressed in < 5 cells were also removed. Cells were classified into muscle, liver, brain, skin, and hematopoietic groups based on tissue type and sorting method. Integration, clustering, and cell annotation were performed for each group as in the SSc. After annotation, all the cells were merged; the top 6,000 highly variable genes were used for PCA and the top 50 principal components were used for UMAP.

### scPAISO pipeline

#### Read1 alignment

For each library, Read1 was aligned to the reference genome (GRCh38 for human, GRCm38 for mouse) using STAR aligner (version 2.7.10b) with the following parameters: *--clip5pNbases 58--outFilterMismatchNmax 999--outFilterMismatchNoverLmax 0.3--outFilterScoreMinOverLread 0--outFilterMatchNminOverLread 0--outFilterMatchNmin 70*. Read2 alignment was performed using the Bam file output by Cell Ranger.

#### Identification of PASs

BAM processing: scanBam (https://github.com/Bioconductor/Rsamtools) was used to read the BAM files, and only paired reads and uniquely mapped reads were retained. Paired reads with a distance > 1 Mb between their mapped positions on the chromosome were filtered. Paired reads were aligned to genes based on Read2 positions; intron lengths were subtracted from Read2–Read1 distances. Paired reads with a distance > 10,000 bp were filtered out.

PAS peak calling: the 5’ ends of Read1 (sense and antisense strands) were extracted separately and exported as BigWig files using the‘export.bw‘function from the rtracklayer package. MACS3 (version 3.0.0)^31^ identified candidate PASs with the following parameters: *--max-gap 10--min-length 5--extsize 3--shift-1--call-summits--keep-dup all-p 0.00001.* Candidate PASs with the 6-mer’AAAAAA’ within 5 bp upstream to 20 bp downstream were filtered to exclude priming artifacts. PASs within 50 bp were further merged into the final PAS.

PAS annotation: PASs were classified as 3’ UTRs, introns, or exons based on their genomic location. The 3’ UTR PASs were further classified as terminal PASs (tPAs), whereas those in introns or exons were classified as intronic PASs (iPAs) or exonic PASs (ePAs), respectively.

#### Model fitting and PA isoform quantification

For the previously described paired Read1 and Read2 sequences, those where Read1 did not overlap with the PAS were excluded, generating a positive control dataset. The intron-removed distance between Read2 and its corresponding PAS was calculated and used to fit a statistical model (Weibull, Gamma, or Log-normal distribution). The fitted model was applied to all Read2 in the positive control datasets to predict the most likely PAS for each read. The model with the highest global accuracy was subsequently used for the quantification of PA isoforms. A single-cell PA isoform count matrix was then generated, where each cell was represented by the expression levels of different PAS isoforms. This matrix was used for downstream analysis, including differential expression, clustering and PAS usage analysis.

#### Rank score calculation

A rank score was calculated for each gene to evaluate the usa of different-length PAS isoforms to quantify PAS usage bias:

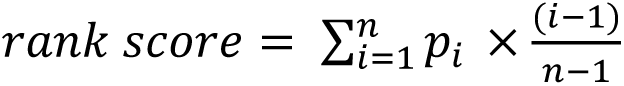

Where *n* denotes the number of PAs, 𝑝_i_ denotes the proportion of expression of the i-th PA relative to the total expression of the gene. The rank score ranges from 0 (exclusive use of the proximal PAS) to 1 (exclusive use of the distal PAS).

### Cell clustering by relative PAS usage

The PA isoform count matrix was processed using PASTA^34^. And CalcPolyAResiduals^34^ was used to calculate the residuals of PA isoforms based on the Dirichlet multinomial distribution, which can characterize heterogeneity in the relative use of PASs at single-cell resolution. FindVariableFeatures^35^ was used to identify polyA sites that varied across cells, after which dimension reduction was performed on the variable polyA sites.

### Differential analysis

For gene and PA isoform-level differential expression analysis, the Seurat package (version 4.0.5) was used. The FindMarkers^35^ function was used to identify differentially expressed genes or PAS isoforms between cell types or conditions (p_val_adj < 0.01 and |avg_log2FC| > 0.3).

Differential relative PAS usage analysis was performed using FindDifferentialPolyA function from PASTA^35^. This function returns the following outputs: *estimate*, estimated coefficient from the differential APA linear model (indicates the magnitude of change); *p.value*, p-value from the linear model, *p_val_adj*, Bonferroni-corrected p-value; *percent.1*, the average percent usage (pseudobulk) of the polyA site in the first group of cells; *percent.2*, the average percent usage (pseudobulk) of the polyA site in the second group of cells. A threshold of *p_val_adj* < 0.05 and a percent change (|*percent.1 - percent.2*|) > 0.05 were used for further filtering.

To compare the overall changes in 3’ UTR length among different conditions, FindDifferentialPolyA was used to identify genes whose relative PAS usage changed. For each gene, we multiplied the estimate by the ordinal number of the PAS and then summed the values. If the result was > 0, it indicated that the overall 3’ UTR length had increased; if < 0, it indicated that the overall 3’ UTR length had decreased.

## Data availability

The single-cell RNA-seq datasets used in this study are available from the Gene Expression Omnibus (GEO) or Genome Sequence Archive (GSA) under the accession numbers GSE196676 (bone marrow hematopoiesis), GSE249279 (SSc), and CRA004660 (mouse tissues). The processed data and analysis results are available upon request.

## Code availability

The scPAISO pipeline, including scripts for data processing, PAS identification, and quantification, is available on GitHub (https://github.com/yongjieliu/scPAISO). Detailed documentation and example datasets are provided to facilitate the use of the proposed pipeline.

## Acknowledgments

This study was supported by funding from the National Natural Science Foundation of China (82273938, 81922068, 82473934, 32100616, 82304511, 82473935); National Key Research and Development Project (2020YFA0113500); Hunan Provincial Natural Science Foundation of China (2023JJ40341, 2023JJ40352); Scientific Research Project of Hunan Provincial Health Commission (A202302077875, B202302078452, China); the Projects of Army Medical University (2022XJS06, 2023XQN20, 2023XQN19, China); the Chongqing Municipal Education Commission Foundation (KJZD-M202412802, China); the Natural Science Foundation of Changsha, Hunan Province (kq2208087, China); and the Key Research and Development Program Project of the State Key Laboratory of Trauma and Chemical Poisoning (2024K004). We gratefully acknowledge the use of publicly available single-cell RNA-seq datasets from the Gene Expression Omnibus (GEO) and the Genome Sequence Archive (GSA). Specifically, data with the accession numbers GSE196676, GSE249279 and CRA004660 were utilized in this study. These datasets provided important resources for method validation and greatly facilitated our research. Writing errors were checked using Grammarly (http://grammarly.com/).

## Author contributions

Yongjie Liu, Youcai Deng, and Yafei Deng conceived and supervised the project. Yongjie Liu and Peiwen Xiong performed the data analyses. Yongjie Liu, Youcai Deng, Yafei Deng, Junping Wang and Yong Zhu interpreted the data and wrote the manuscript. Songyang Li, Xinjia Liu, Tao Liu, Qinglan Yang, Shuting Wu, Hongyan Peng, Yana Li, and Lingling Zhang reviewed and revised the manuscript.

