## Supplementary Figure S1-7 for "Identification and quantification of alternative polyadenylation sites in single cell RNA-seq data using scPAISO"

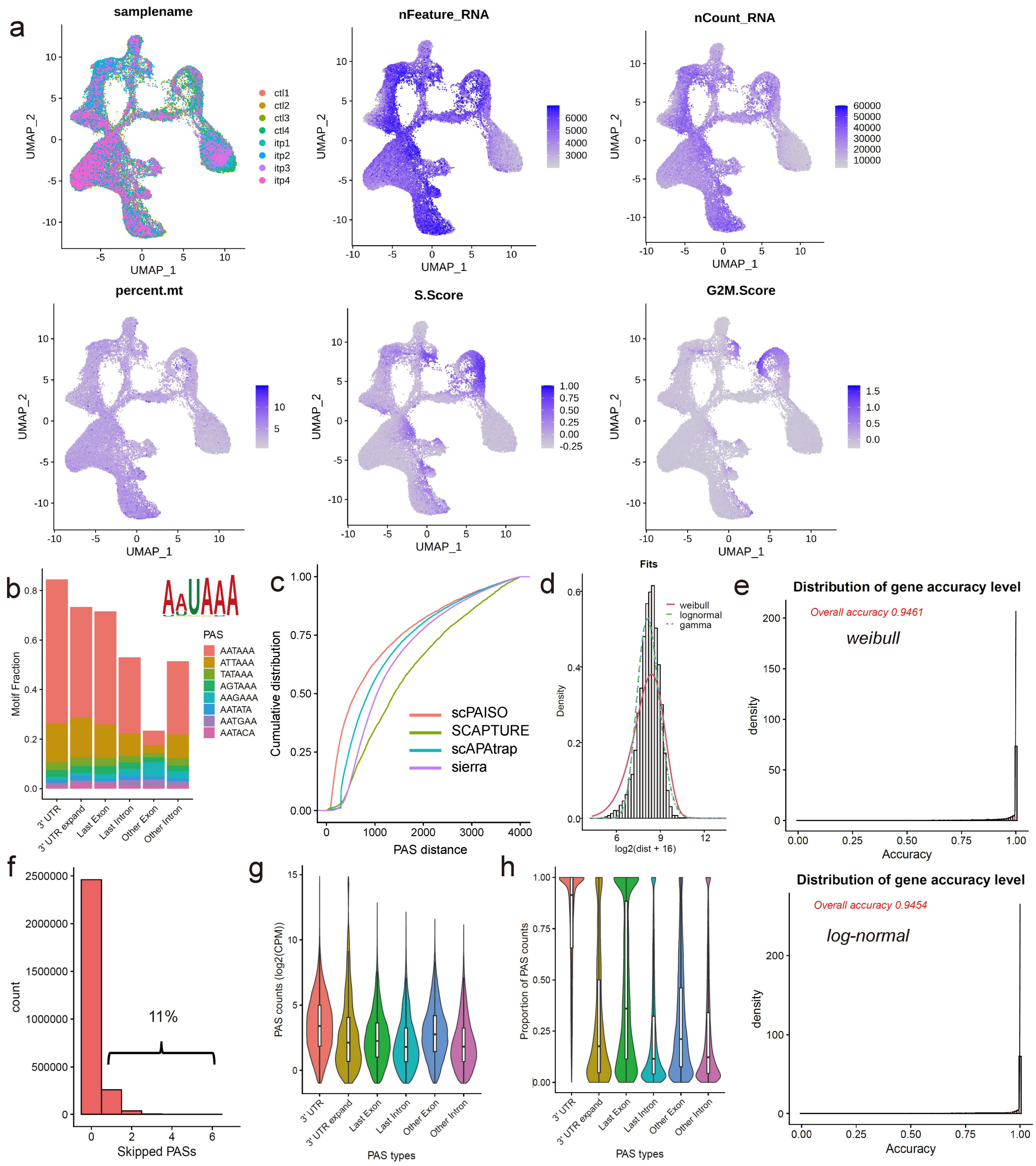


**Supplementary Fig. 1: QC of gene expression and PAS assignment.** (**a**) UMAP visualization of samples, gene numbers, UMIs, mitochondrial counts, S phase scores, and G2/M phase scores. (**b**) Enrichment of the AAUAAA motif in PAS peaks. (**c**) Cumulative distribution of distances between nearby PASs. (**d**) Distribution of distances between the 3' end of Read2 and corresponding 5' end of Read1 (left) or the corresponding PAS (right) in the positive control dataset. (**e**) Model fitting accuracy for the distance between Read2 and PAS using *Weibull* and *log-normal distributions*. (**f**) Bar plot showing the distribution of the number of PASs between Read2 and its corresponding PAS. Most Read2 use the nearest PAS. (**g-h**) Expression levels of PA isoforms (**g**) or relative PA usages (**h**) across different genomic regions.

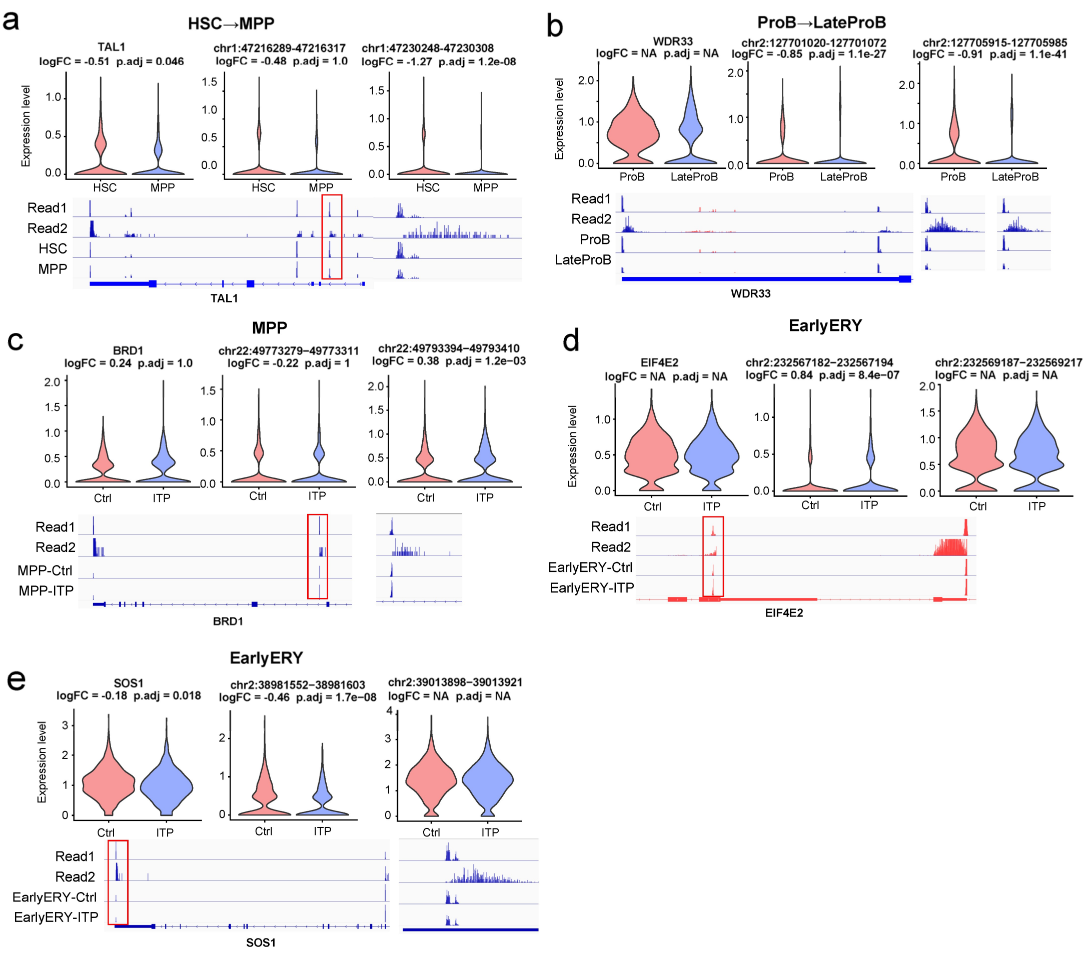


**Supplementary Fig. 2: Changes in expression of the PA isoform.** (**a**) An iPA of *TAL1* showed decreased expression between HSCs and MPPs. (**b**) Two tPAs of *WDR33* showed decreased expression between ProB and LateProB. (**c**) An iPA of *BRD1* showed increased expression in MPP between healthy donors and ITP patients. (**d-e**) An ePA of *EIF4E2* showed increased expression in EarlyERYs between healthy donors and patients with ITP (**d**), whereas a tPA of *SOS1* showed decreased expression (**e**).

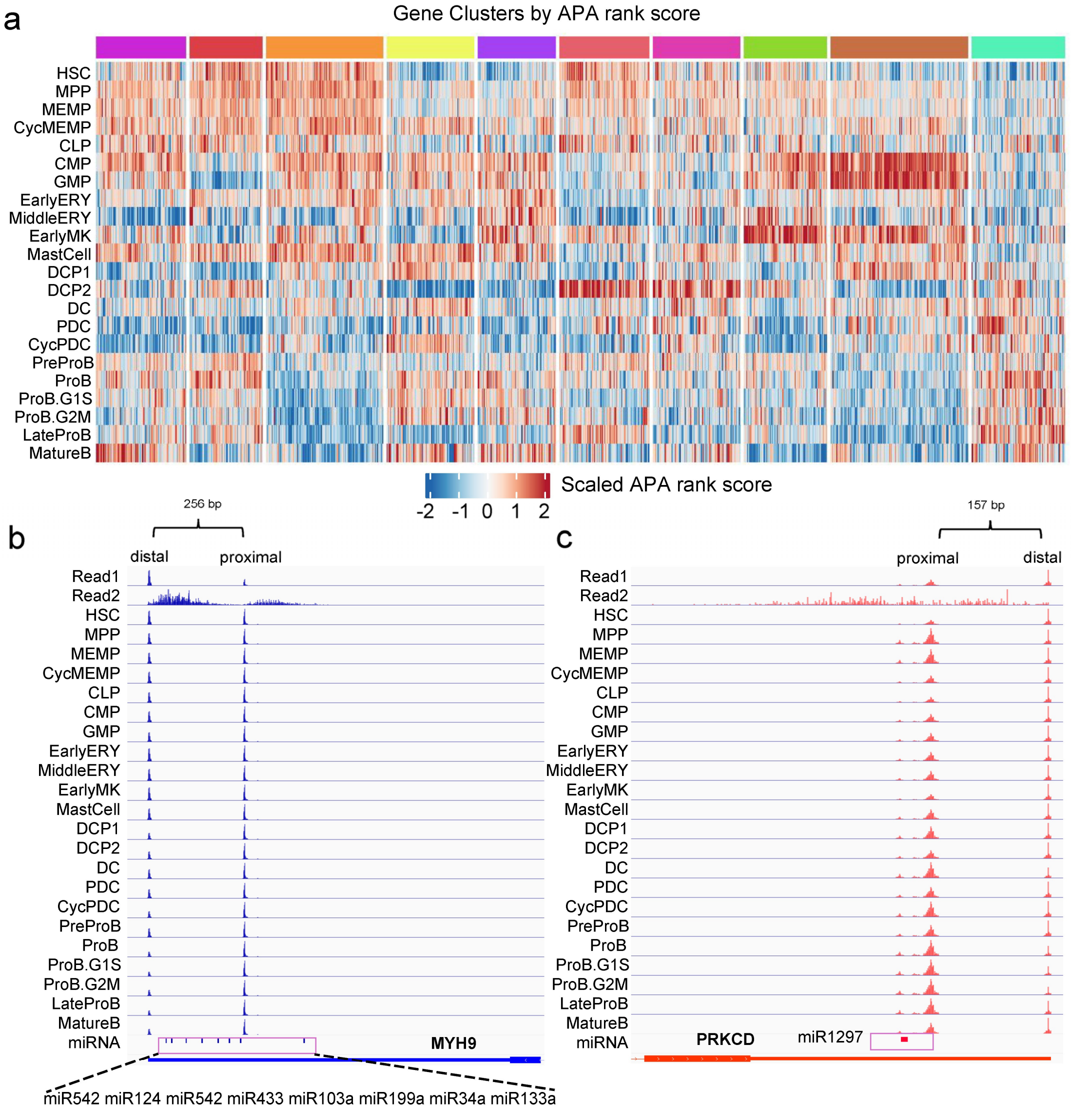


**Supplementary Fig. 3: APA rank score changes in hematopoiesis.** (**a**) Heatmap showing the dynamic normalized rank score across different cells. (**b-c**) Track view of the PAS usage of *MYH9* (**b**) and *PRKCD* (**c**).

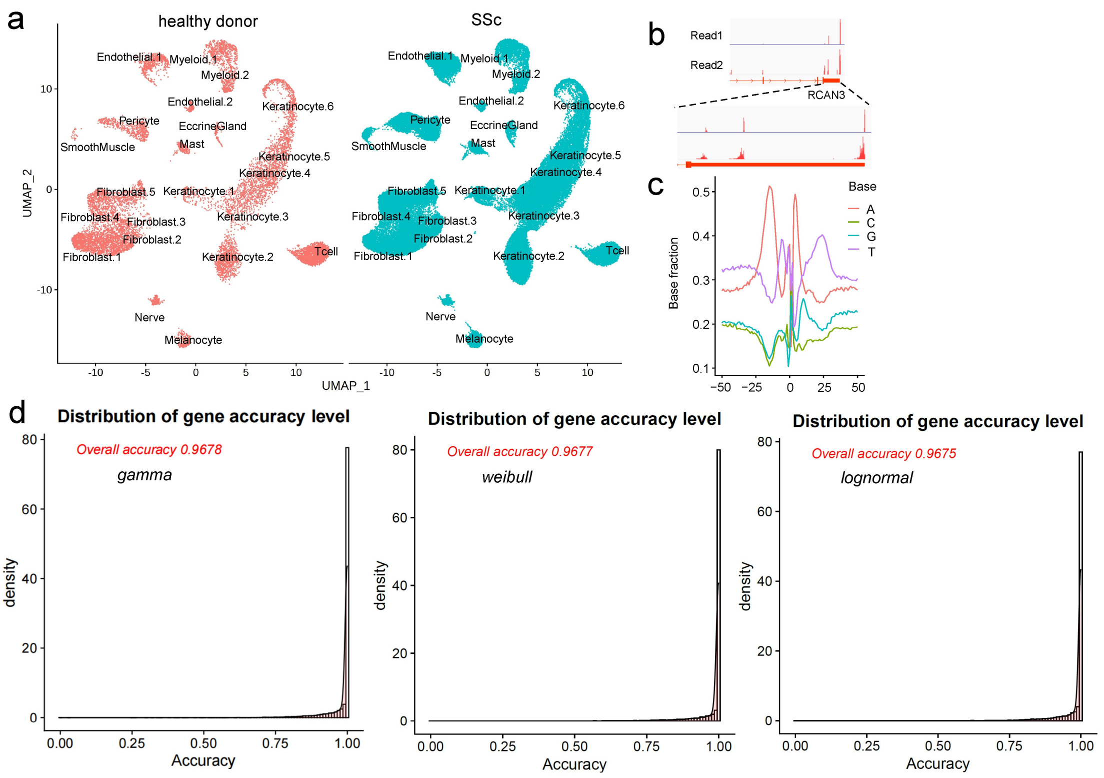


**Supplementary Fig. 4: QC of UMAP visualization and model fitting accuracy in the SSc dataset.** (**a**) UMAP visualization of scRNA-seq data seprated by healthy donors and patients with SSc. (**b**) Example of identified PASs in the 3' UTR of *RCAN3*. (**c**) Nucleotide composition surrounding PASs in SSc, which are consistent with known polyadenylation signals. (**d**) Model fitting accuracy for distances between Read2 and PASs using *gamma*, *Weibull* and *log-normal distributions*.

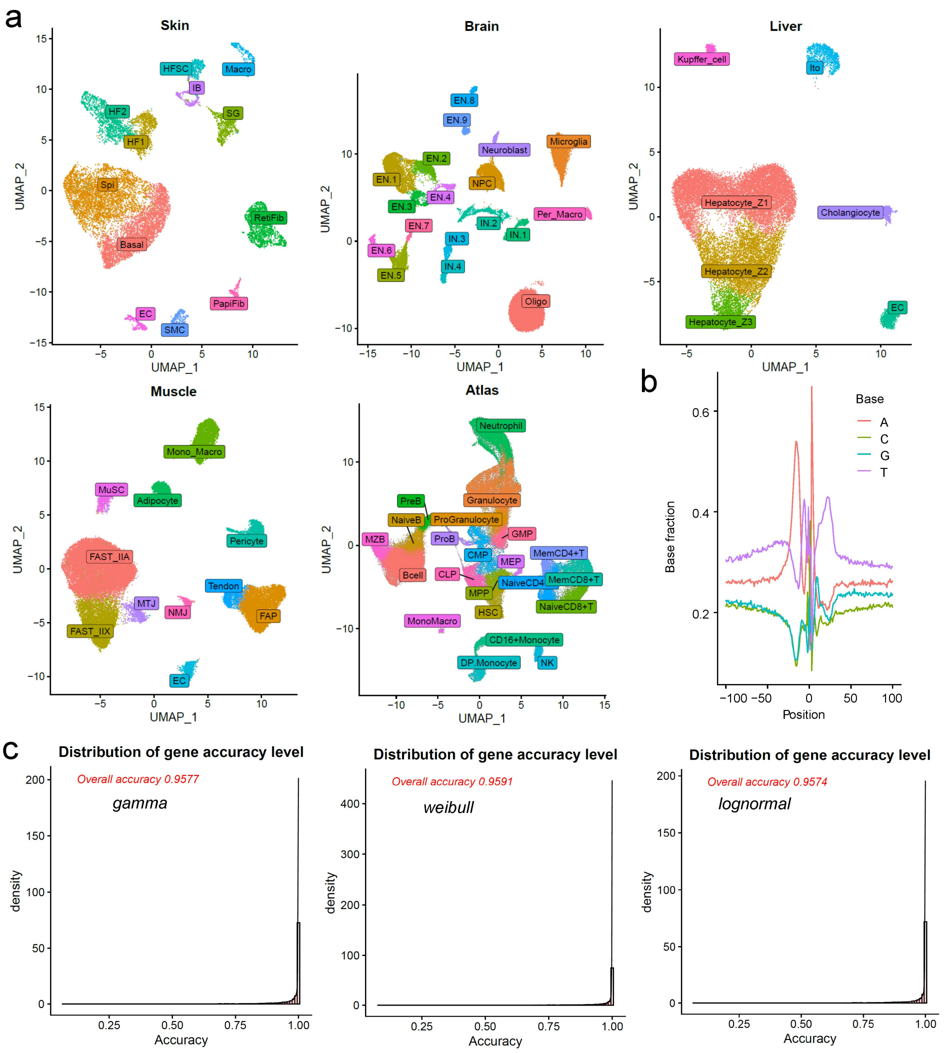


**Supplementary Fig. 5: UMAP visualization and model fitting accuracy in mouse tissue datasets.** (**a**) UMAP visualization of scRNA-seq data from the skin, brain, liver, skeletal muscle and hematopoietic/immune system. (**b**) Nucleotide composition surrounding PASs in mouse tissues. (**c**) Overall accuracy of PAS prediction using the *gamma*, *Weibull* and *lognormal distribution* models.

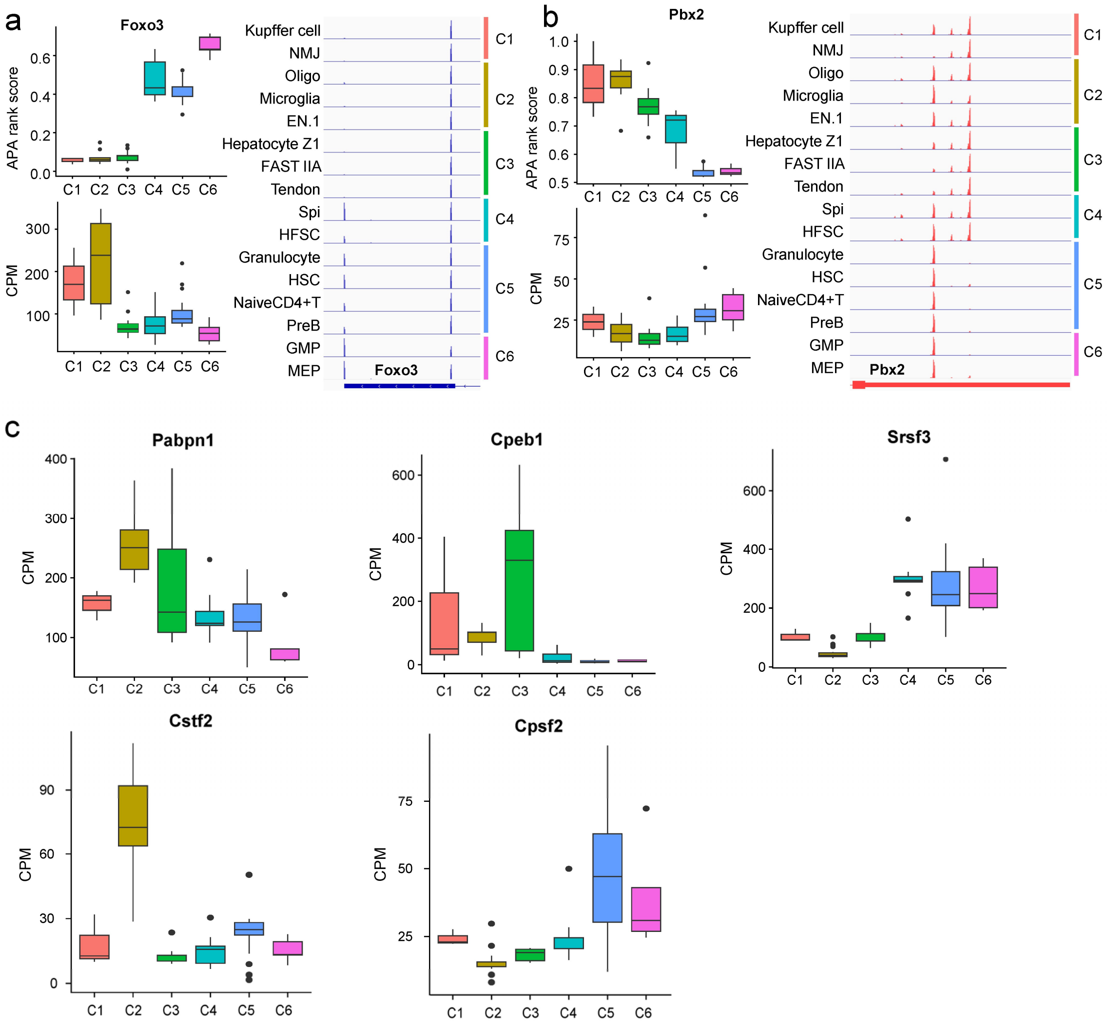


**Supplementary Fig. 6: Gene expression patterns of mRNA 3' end processing factors.** (**a-b**) Tissue-specific PAS usage preferences of *FOXO3* (**a**) and *PBX2* (**b**) across cell clusters. (b) Gene expression preferences of *Pabpn1*, *Cpeb1*, *Cstf2*, *Cpsf2*, and *Srsf3* across cell clusters.

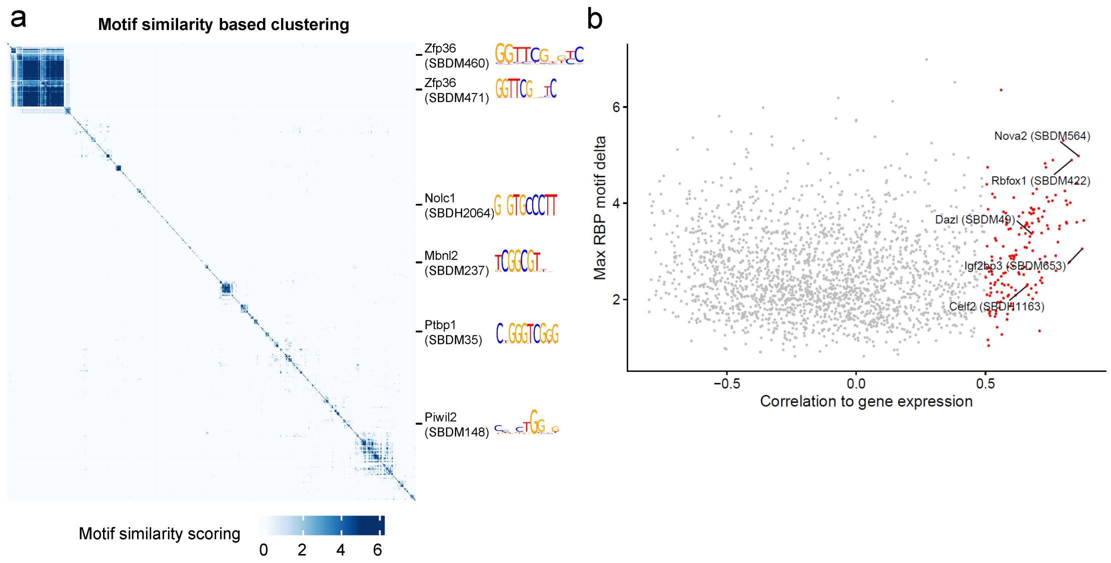


**Supplementary Fig. 7: Motif enrichment analysis of RBPs.** (**a**) Clustered heatmap showing the similarity scores of 2,179 motifs from 319 RBPs calculated by tomtom. (**b**) Scatter plot showing the correlations between gene expression and motif enrichment scores for RBP-motif pairs, with 169 pairs (red points) exhibiting Pearson correlations greater than 0.5.
